# Variational Autoencoder-enabled High-throughput Drug Screening for HIV Latency Modulators predicted through Noise in Gene Expression

**DOI:** 10.64898/2026.07.08.737074

**Authors:** Yiyang Lu, Jesse Horne, Xuenan Mi, Shivank Nag, Sohom Dash, Roy D. Dar, Diwakar Shukla

## Abstract

Due to its ability to establish a pool of undetectable and latently infected cells that can initiate viral production through random reactivation, a cure to human immunodeficiency virus (HIV) infections has remained elusive. Many approaches have been proposed, including the “shock and kill” method where latency reversing agents (LRAs) are administered to reactivate latently infected cells out of latency and remove them through immune targeting and clearance, and the “block and lock” method where latency promoting agents (LPAs) are administered to inhibit reactivation and potentially induce a “deep latency” state where infected cells can no longer reactivate. Previous large scale drug screen studies have demonstrated a correlation between a compound’s capability to modulate the fluctuations (or “noise”) in HIV gene expression and its potential to modulate HIV latency. However, measurements of gene expression noise are labor-and cost-intensive. To circumvent these drawbacks, we trained a variational autoencoder (VAE) on a previously published large scale time-lapse fluorescence microscopy dataset, and performed an *in silico* screening of ∼175,000 compounds for HIV latency modulators. Out of the top 113 predicted modulators that were experimentally tested, 16 latency reversing agent (LRA) synergizers and 2 latency promoting agents (LPAs) were confirmed, yielding an overall experimental hit rate of 15.9%. Our work demonstrates that *in silico* drug screening modalities, guided by existing large-scale experimental datasets, can yield high experimental hit rates, reducing costs incurred from labor-intensive wet lab-focused methodologies.

## Introduction

Human immunodeficiency virus (HIV) belongs to the genus *lentivirus* of the *retroviridae* family. Without treatment, HIV infection develops into acquired immunodeficiency syndrome (AIDS)^1^, where the patient’s immune system is severely weakened and opportunistic infections become life-threatening^2^. Since the initial discovery of HIV in 1983^3^, enormous amount of research has been dedicated to treating and curing HIV infection. However, cells infected by HIV can enter a prolonged and inactive state called latency, in which they cannot be targeted by either drug treatments or the immune system. These cells can be maintained in the body for decades^4,5^, and decay extremely slowly (half life of ∼44 months)^6^. Thus, methods to eliminate HIV from patients have remained elusive despite continued efforts from the research community. While antiretroviral therapies (ARTs) have been proven highly effective at controlling viral loads, patients need to remain on ART for the rest of their lives. This is because without constant ART administration, latently infected cells can spontaneously and stochastically initiate viral production, through a process termed reactivation. This eventually results in a rebound in viral load and accelerates the infection towards AIDS.

Many strategies have been devised to tackle HIV latency. Two of the most prominent strategies are termed “Shock and Kill” and “Block and Lock”^7^. The “Shock and Kill” strategy aims to fully reactivate all latently infected cells (“Shock”) by using latency reversing agents (LRAs) and remove them from the patient through immune targeting and clearance (“Kill”). The “Block and Lock” strategy aims to inhibit and stop reactivations (“Block”) by using latency promoting agents (LPAs) and induce a “deep latency” state where the latently infected cell can no longer spontaneously reactivate (“Lock”). While “Shock and Kill” has been a more prominent research topic than “Block and Lock”, the effectiveness and safety of existing compounds within the repertoires of both strategies continue to be underevaluated^8,9^. Many promising compounds that reactivate HIV *in vitro* either failed to generate similar levels of reactivation *in vivo*, or exhibit off-target effects^10,11^. On the other hand, compounds capable of “Block and Lock” have been difficult to come by, and current ongoing research has yielded few notable results^12^.

The integrated HIV provirus is known to exhibit strong fluctuations in its gene expression (or “noise”), which has been indicated as a key player in the decision-making of latency versus active replication^13^. The HIV long terminal repeat (LTR) promoter contributes to the noise by generating random bursts of transcripts among long periods of no transcriptional activity^14^. The HIV transactivator of transcription (tat) protein also contributes to the overall noise by providing a strong positive feedback loop to its transcription after significant accumulations. This results in bifurcating phenotypes within a single clonal (i.e. cells with identical genetic makeups) population, where at any given time, some cells exhibit high levels of HIV expression while others maintain a minimal basal expression level close to zero^15^. Gene expression noise has been utilized as a drug screening criterion in the search of chemical compounds capable of modulating HIV gene expression. Among these compounds, many were discovered to be capable of modulating HIV latency as well, including some that synergize with existing latency reversing agents (LRA synergizers), and some that promote HIV latency (LPAs)^16,17^.

In a previous study by Lu et al^17^, through the combination of a time-lapse fluorescence microscopy primary screen for HIV noise modulators, and a flow cytometry secondary screen for HIV latency modulators, a correlation was discovered between the two types of modulators. However, time-lapse fluorescence microscopy is more time-and resource-intensive compared to flow cytometry, and its use as a primary screen did not provide an increase in throughput compared to direct screening of the entire library with flow cytometry-based assay. On the other hand, machine learning techniques, specifically deep learning, have been widely used to predict molecular properties and provide efficient identification of drug candidates to treat many infectious diseases. Given the large experimental dataset generate from Lu et al^17^ containing the time-dependent response of HIV gene expression to treatments of various small molecules, deep learning methodologies are well suited to extract meaningful information from this dataset containing responses of HIV to known LPAs and LRA synergizers, and enable the identification of novel HIV latency modulators. Advanced deep learning models, such as variational autoencoders (VAE)^18^ and generative adversarial networks (GAN)^19^, have been developed to extract low-dimensional representations from high-dimensional time-series data. VAE-based methods have been applied to analyze single-cell RNA sequencing data without the need for data preprocessing^20^, while GAN-based frameworks have demonstrated superior predictive abilities compared to conventional models by generating realistic time-series data for stock market prices, appliance energy usage in buildings, and lung cancer progression^21^.

In this study, we aim to utilize the large scale time-lapse fluorescence microscopy data generated in Lu et al^17^ that captures HIV gene expression noise in host cells when exposed to a variety of compounds, to train a machine learning model for the prediction of novel compounds capable of modulating HIV latency (Fig. 1, A, B and C). Using unsupervised learning, we mapped compounds to a latent space, enabling the identification of unique signatures associated with LPAs and LRA synergizers. We integrated chemical structure information with time-lapse fluorescence microscopy data to train our generative model based on the variational autoencoder (VAE) architecture. We tested various encoding strategies to embed the chemical structure information of drug compounds, including transformer-based encoders and continuous latent space encoding^22^. From a university compound library containing about 175,000 compounds^23^, we identified potential LPAs and LRA synergizers in the latent space that closely resembled known LPAs and LRA synergizers. Out of the top 113 predicted compounds, 16 LRA synergizers and 2 LPAs were validated experimentally, yielding an experimental hit rate of 15.9%. This is a significant improvement over existing high-throughput drug screening modalities, which frequently result in experimental hit rates of no higher than 2%^24–26^. Taken together, our study presents a novel computational workflow for identifying HIV latency modulators by capturing distinctive patterns in the VAE latent space, providing an alternative primary screen modality to time-lapse microscopy with massively improved throughput. The results from our computational workflow were experimentally verified on HIV latency model cell lines covering multiple integration sites, and demonstrated enhanced experimental hit rates in the HIV drug screening process.

**Figure 1.**
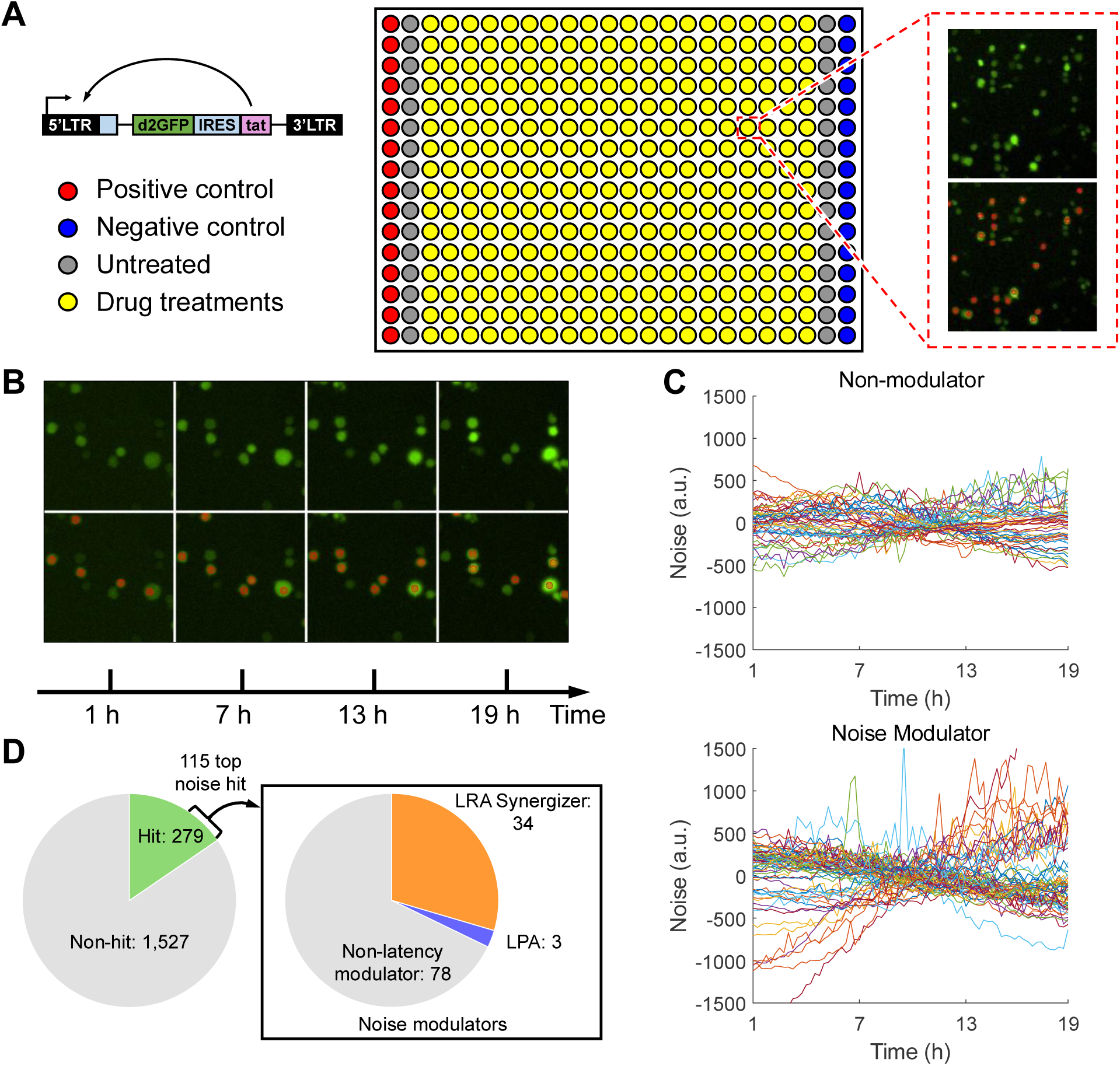
Source of the training dataset of the ML algorithm. The training dataset was experimentally collected by Lu et al., PNAS 2021^17^. **(A)** Experimental setup for the time series data. Jurkat cells containing the Ld2GIT construct, a minimal HIV feedback gene circuit (left), were plated in 384-well plates, and treated to various conditions, including 10 ng/ml of TNF (positive control), 2 μM of Alp (negative control), untreated (neutral control), and 10 μM of one of the 1,806 small molecules from a diverse library (middle). The cells were continuously imaged for 48 hours, with one image taken for each well every 15 minutes. Each image was segmented by a binary mask to identify individual cells (right). **(B)** An example of the resulting time-series images. Images from hours 1 – 19 were further analyzed to allow the cells sufficient time to adapt to the plate environment, and to avoid cell detachment that happens after 20 – 24 hours of imaging. **(C)** Example noise trajectories from a non-modulator compound (top) and a noise modulator compound (bottom) quantified through microscopy image analysis. **(D)** The initial fluorescence microscopy-based screening resulted in 279 hit compounds that significantly modulated fluctuations in the expression of the Ld2GIT construct (left). A secondary screen on HIV latency cell models on the 115 top hits revealed 34 LRA synergizers and 3 LPAs (right).

## Results

### Variational Autoencoder Identifies HIV Gene Expression Modulators

We utilized a large-scale HIV drug screening dataset utilizing time-lapse fluorescence microscopy and flow cytometry published by Lu et al^17^. The microscopy dataset consists of 18-hour single cell time-lapse fluorescence microscopy data, curated from multiple 48-hour time-lapse experiments using the Ld2GIT HIV model cell line. The 18-hour data consists of images taken between hours 1 – 19 of the 48-hour experiments to account for the initial adaptation period after adhering the cells to the imaging plates, as well as to exclude excess cell movements and detachments after 20 hours. The Ld2GIT cell line consists of Jurkat cells, each containing a stably integrated minimal HIV feedback gene circuit (Fig. 1A, left). The gene circuit consists of the HIV 5’ LTR promoter, followed by a destabilized GFP element (d2GFP, protein half-life = 2.55 h), an internal ribosome entry site (IRES), and the HIV transactivator of transcription (tat)^27,28^. The tat protein, once expressed, enhances the transcriptional efficiency of the LTR promoter, forming a positive feedback loop. The cells were adhered to the glass bottom of 384-well plates using cell adhesive, and each well was treated with one of the following conditions: 10 ng/ml of tumor necrosis factor (TNF, positive control, enhances HIV expression), 2 μM of alsterpaullone (Alp, negative control, suppresses HIV expression), untreated (neutral control), or 10 μM of one of the 1,806 compounds from a diverse library of chemicals (Fig. 1A, middle)^17^. The plate was imaged on an inverted fluorescent microscope to monitor the accumulation of d2GFP inside the cells for a total duration of 48 hours, and the resulting images were segmented into binary masks of single cells (Fig. 1A, right) which were tracked through the time series (Fig. 1B). The time-lapse fluorescence microscopy experiment identified a total of 279 initial hit compounds that significantly modulated the fluctuations (or “noise”) in the expression of the Ld2GIT vector (Fig. 1C, left). The top 115 hits were then examined with a latent HIV cell model JLat. Out of the 115, 34 compounds synergized with existing latency reversing agents (LRAs) to enhance latent HIV reactivation (Fig. 1C, right), classifying them as LRA synergizers. Another 3 compounds suppressed HIV reactivation when administered together with known LRAs, indicating that they belong to the category of latency promoting agents (LPAs) (Fig. 1C, right). Due to the imbalance between the available microscopy data for HIV latency modulators (LPAs and LRA synergizers) and non-modulators, an additional 12 time-lapse fluorescence microscopy measurements were conducted on 4 previously reported LPAs (PX12, tiopronin, NSC 400938, NSC 155703), as well as TNF and untreated conditions as controls, under the same protocol from Lu et al^17^.

We applied a generative model based on the variational autoencoder (VAE) architecture^18^ to capture latent representations from 1,838 independent microscopy measurements of single-cell time-series protein expression for 1,806 compounds treated as chemical perturbations (see Materials and Methods for details on the quality control of microscopy data). The molecular structures of the compounds and their 18-hour fluorescence noise trajectories were used as input to the VAE to infer their hidden representations in a low-dimensional space. After testing various encoder models including MoLFormer^29^ and Mol2vec^30^, molecular structure information was finally extracted with CDDD (Continuous Data-Driven Descriptors), which uses a neural machine translation framework translating between chemical string representations^22^ (Fig S1). The model was pretrained on a large dataset of SMILES (Simplified Molecular Input Line Entry System)^31^ from the ZINC and PubChem databases, yielding fixed-length, continuous embedding vectors^32,33^. We applied average pooling over atomic tokens for each molecule to generate a 512-dimensional vector representing the molecule’s structural embeddings. These molecular embeddings were then concatenated with time-series fluorescence noise trajectories to serve as the input to the VAE model, allowing it to learn the latent space representation for each compound in the experiment over many training epochs. We systematically ablated our model architectures to test conditional (CVAE), multi-modal (MMVAE), hierarchical (HHVAE), and X-shaped (XVAE)^34^ variational autoencoders with various latent space sizes and embeddings (Fig. S1). Our loss-optimized XVAE model consists of two hidden layers of 256 and 32 neurons, and is parametrized with a latent dimension of 4, a batch size of 128, and a learning rate of 0.001. The model completes 95% of its learning by the 256^th^ epoch, while convergence is achieved by epoch 380, beyond which the marginal improvements in validation loss become negligibleAfter 100 training epochs, the VAE model achieved convergence based on both training and validation loss. (Fig. S2).

The trained VAE model was then applied to screen for potential LPA and LRA synergizers within a collection of chemical libraries hosted by the High Throughput Screening Facility (HTSF) at University of Illinois Urbana-Champaign, including the Chembridge MicroFormat library, the HTSF library, the Marvel library, and four libraries from the National Cancer Institute (NCI): Diversity set, Natural Products Set, Challenge Set, and Open Plate Set^23^. As shown in Figure 2A, the workflow comprised of training and inference stages. For screening, we processed approximately 175,000 compounds from HTSF, using CDDD to extract their chemical embeddings and concatenating them with latency modulator noise trajectories from Lu et al^17^ to infer their likelihood to modulate HIV latency. The concatenated vectors were then passed through the VAE to map each compound onto the latent space. By calculating the pairwise distances between queried compounds and known LPA and LRA synergizer compounds, we ranked the candidates by similarity, selecting the top 113 latency modulators for further experimental validation. Figure 2B shows the VAE latent space of 1,838 individual time-lapse fluorescence microscopy measurements, highlighting distinct clustering patterns across different categories. Known activators (TNF-treated, positive control), known suppressors (Alp-treated, negative control), and untreated measurements each form well-defined clusters that are clearly separated from one another.

**Figure 2.**
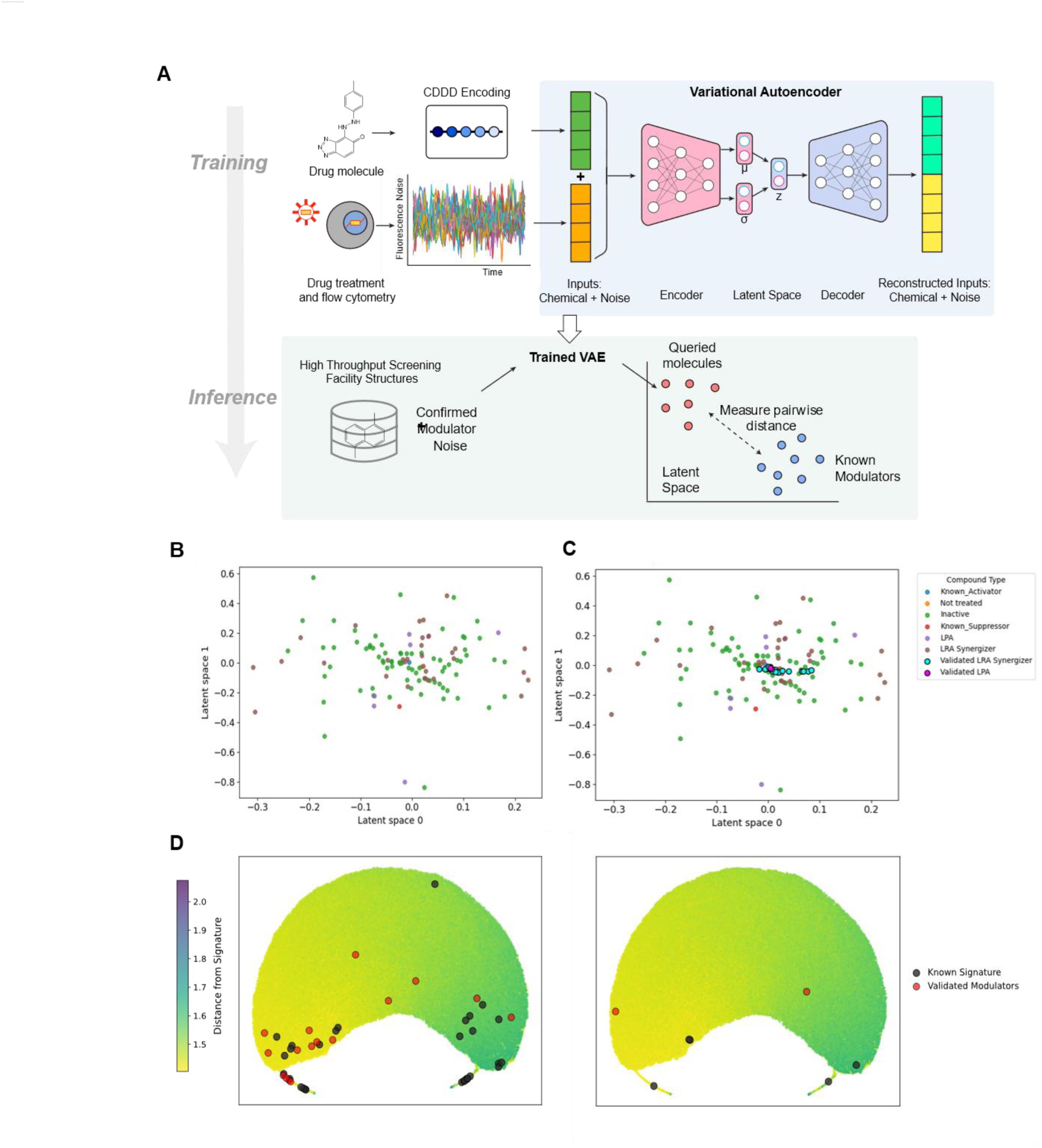
Workflow of identifying HIV latency modulators. **(A)** The workflow consists of training and inference stages to screen for potential modulators of HIV gene expression. In the training stage, single-cell time-series protein expression data and molecular structure embeddings, generated using CDDD, are combined as inputs to a VAE model to learn their hidden representations. During inference, the trained VAE model is used to analyze a large compound library by comparing latent space distances between candidate compounds and known modulators, selecting the top candidates for experimental validation. (**B**) VAE latent space of 1,838 independent measurements with single-cell time-series protein expression data. (**C**) Projection of the two experimentally validated LPAs and sixteen LRA synergizers onto the trained VAE latent space. **(D)** UMAP projection of validated and known LRA synergizers (left) and LPAs (right) onto background test molecules colored based on distances from the respective known signatures.

### ML Predictions Demonstrate High Experimental Hit Rate

To validate the predictions generated by our ML algorithm, we utilized the established method of latency reversal assay to examine whether the predicted compounds can modulate HIV latency. Top-ranked predicted compounds were challenged with the known LRA tumor necrosis factor (TNF) to probe their interactions with it. To that end, cultures of a previously reported HIV latency model cell line, JLat 9.2^35^, were treated with each predicted compound separately. JLat 9.2 cells contain the full-length HIV-1 genome, with a frameshift mutation of the *env* reading frame, and a deletion of the *nef* reading frame. A DNA sequence encoding green fluorescent protein (GFP) is inserted in place of *nef* ^35^ (Fig. 3A). For each compound, two conditions were tested: with and without the addition of TNF (Fig. 3B). This allows for the quantification of the interaction between the compounds and TNF using the Bliss independence model^36^. ML-predicted compounds were added at a final concentration of 5 μM, while TNF was added at 10 ng/ml. Cells were treated for 24 hours at 37 °C and 5% CO_2_ incubation conditions, and measured on flow cytometers to quantify cell viability and latent HIV reactivation (Fig. 3B, 3C and 3D, see Fig. S3 for gating strategy). An excess over Bliss (EoB) score is calculated for each predicted compound (Methods), where a positive score indicates synergy between the compound and TNF, and a negative score indicates antagonism. The magnitude of the score indicates the strength of the interaction. Due to throughput constraints, the validation assay was carried out on 5 separate days (Fig. 3D and 3E, days separated by vertical dashed lines). To reduce false positives, each treatment condition was carried out in duplicates (n = 2), and a median threshold was applied to the magnitude of EoBs for each day of experiment (Fig. 3E, horizontal dashed lines). Among the top 113 predicted candidates, 2 LPAs (antagonize TNF and suppress reactivation of latent HIV) and 16 LRA synergizers (synergize with TNF to reactivate latent HIV) were discovered (Fig. 3C, 3D and 3E, see Fig. S4 for full result), yielding an experimental hit rate of 15.9%. The percentage of cells reactivated from the latency reversal assay showed weak correlation with the normalized EoB score of each compound (Fig. 3C). This is due to the EoB score normalizing the reactivation percentage according to the TNF-only control samples on each day. Two compounds yielded high reactivation percentages but were not classified as LRA synergizers, due to either high reactivation without TNF treatment (compound A13, Fig. 3C, Fig. S3A) or cytotoxicity (compound A15, Fig. 3C, Fig. S3A). Six other compounds suppressed reactivation percentages below TNF control and resulted in significantly lowered EoB scores, but were not classified as LPAs due to cytotoxicity (compounds B13, D13, E15, E17, F15, G15, Fig. 3C, Fig. S3A, B and C). A summary of the hit compounds can be found in Table 1. Figure 2C shows the projection of two experimentally validated LPA molecules and sixteen verified LRA synergizers onto the trained VAE latent space. The sixteen LRA synergizers are positioned relatively close to each other, and while there is some overlap with LPA molecules, it could be possible that the two classes are more distinguishable in the higher-dimensional latent space.

**Figure 3.**
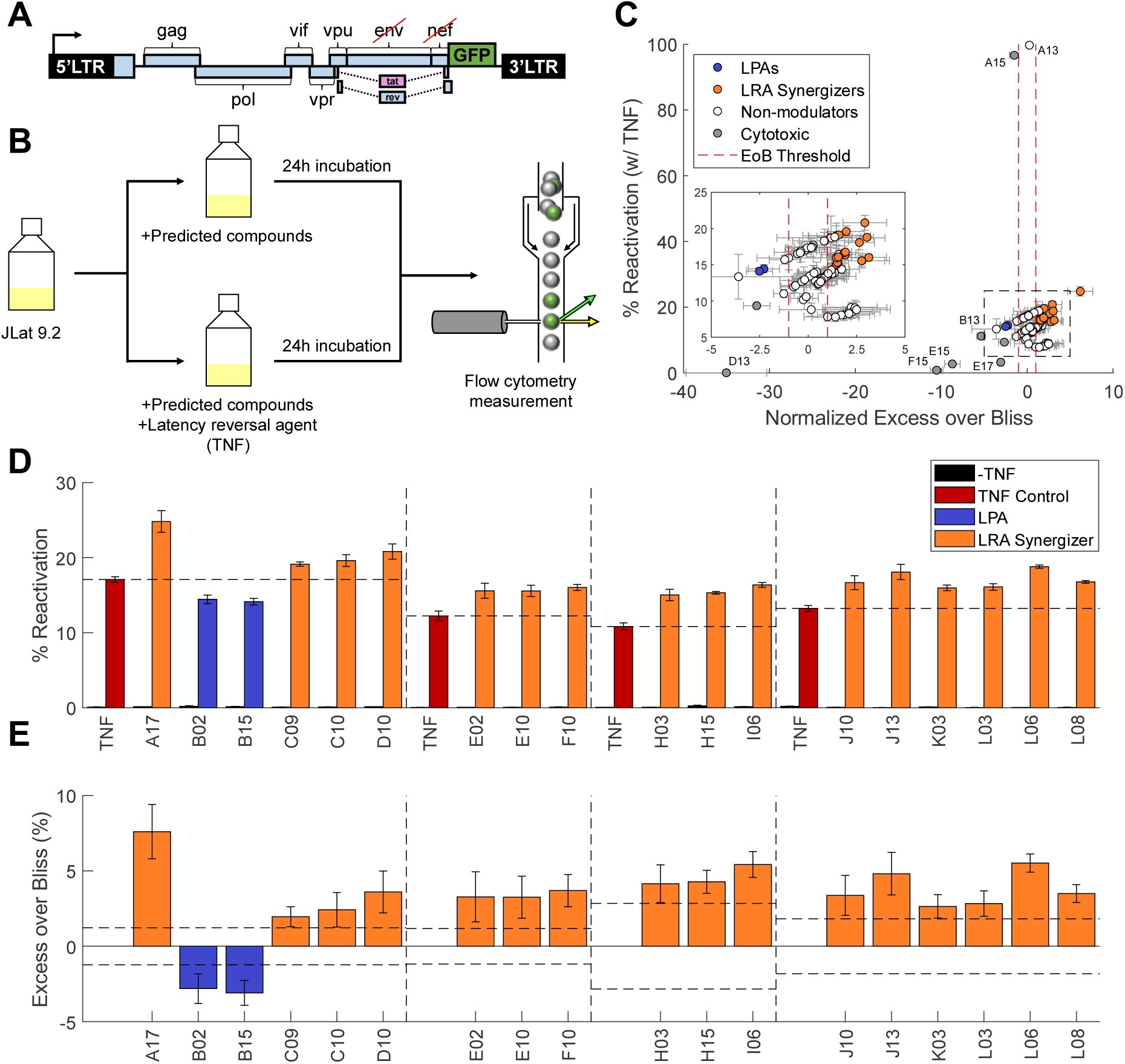
Experimental validation of 113 ML-predicted molecules yielded 18 latency-modulating compounds. **(A)** The HIV latency cell model, JLat, contains the HIV-R7/E^-^/GFP construct^35^. It is based on the full-length HIV-1 genome, with a frameshift mutation in the env reading frame, and a deletion of the nef reading frame, replaced with a GFP element. **(B)** The experimental workflow of the latency reversal assays. JLat 9.2 cell culture were treated with a predicted compound, with and without the addition of TNF. After 24 hours of incubation at 37 °C and 5% CO_2_, the samples were measured on a flow cytometer. **(C)** Scatter plot representation of the reactivation percentage for each of the 113 tested compounds and their corresponding excess over Bliss (EoB) score, normalized to the EoB threshold for each day of experiment. Treatments were done on JLat 9.2, with each sample treated with one of the 113 tested compounds plus TNF for 24 h. Inset shows a zoom-in view for the cluster of compounds within the black dashed lines. Red vertical lines indicate the thresholds for a compound to be considered an LRA synergizer or an LPA. See panel (E) for more details. **(D)** The percentage of cells that reactivated after drug treatments. TNF: tumor necrosis factor alpha. Other x-axis labels correspond to compounds predicted by ML. For each pair of bars, the left bar (black) represents when the cells were only treated with the labeled compounds, while the right bar (orange/blue, colored according to their interactions with TNF) represents when the cells were treated by both the labeled compounds and TNF. Horizontal dashed lines indicate reactivation percentages from the TNF-only controls. Vertical dashed lines separate different days of experiments. Concentrations: ML predictions: 5 μM; TNF: 10 ng/ml **(E)** The excess over Bliss (EoB) score of each compound, in percentage. Horizontal dashed lines indicate the EoB score thresholds for each drug to be considered as a LPA or LRA synergizer. Vertical dashed lines separate different days of experiments. Only hit compounds are shown in this figure to reduce clutter. Experiment on day 5 did not yield any hits. See Figure S3 for gating strategy and Figure S4 for the full results.

**Table 1.** List of latency modulating compounds. The table includes details of the 18 latency modulating compounds validated during the JLat 9.2 latency reversal assay. Sources of compounds are included. The validation results on the three JLat cell lines are summarized here as well. Testing results labeled as N/A in the JLat 8.4 & 6.3 column are due to the compounds being unavailable from the supplier. NCI: National Cancer Institute. Compounds from ChemBridge can be ordered from Hit2Lead.com. UIUC: University of Illinois Urbana-Champaign. Compounds from UIUC can be requested from the High-throughput Screening Facility (HTSF library: https://scs.illinois.edu/htsf-library; Marvel library: https://scs.illinois.edu/marvel-library)

| Compound | Source | ID | JLat 9.2 | JLat 8.4 & 6.3 |
| --- | --- | --- | --- | --- |
| A17 | NCI | NSC 90786 | LRA Synergizer | LRA Synergizer |
| B02 | UIUC | HTSF 14B19 | LPA | N/A |
| B15 | NCI | NSC 345647 | LPA | (cytotoxic) |
| C09 | ChemBridge | 5552693 | LRA Synergizer | LRA Synergizer |
| C10 | ChemBridge | 6152241 | LRA Synergizer | LRA Synergizer |
| D10 | ChemBridge | 5477479 | LRA Synergizer | LRA Synergizer |
| E02 | UIUC | HTSF 14M22 | LRA Synergizer | N/A |
| E10 | ChemBridge | 7488842 | LRA Synergizer | LRA Synergizer |
| F10 | ChemBridge | 7447396 | LRA Synergizer | LRA Synergizer |
| H03 | UIUC | HTSF 2H10 | LRA Synergizer | LRA Synergizer |
| H15 | NCI | NSC 36572 | LRA Synergizer | LRA Synergizer |
| I06 | UIUC | Marvel 4C21 | LRA Synergizer | LRA Synergizer |
| J10 | ChemBridge | 7664353 | LRA Synergizer | LRA Synergizer |
| J13 | NCI | NSC 307241 | LRA Synergizer | LRA Synergizer |
| K03 | UIUC | HTSF 2L9 | LRA Synergizer | LRA Synergizer |
| L03 | UIUC | HTSF 2M9 | LRA Synergizer | LRA Synergizer |
| L06 | UIUC | Marvel 5F3 | LRA Synergizer | LRA Synergizer |
| L08 | ChemBridge | 5675891 | LRA Synergizer | LRA Synergizer |

Figure 2D shows the validated candidates in close proximity to previously known modulators in the low distance neighborhoods of a UMAP-projected ∼170,000 test molecule background space.

### Investigation of Integration Site Dependency of the Hit Compounds

Belonging to the genus of lentivirus, HIV is capable of stably integrating its viral DNA into the host cell genome. The integration process is known to be semi-random, with preferences towards loci containing actively transcribed genes to promote efficient transcription^37,38^. The result of such behavior is a vast array of integration sites scattered across all 23 pairs of chromosomes, with the integration site densities varying in accordance with the density of endogenous human genes^37^. In addition, previous literature has reported extensive evidence that the expression level of integrated HIV provirus is highly dependent on its neighboring genetic landscape. Level of transcription activities, chromatin structures, transcriptional orientation of the provirus, and the specific genes surrounding the integration site are among the genetic features that influences HIV expression^35,39–42^. However, the JLat 9.2 cell line is a clonal cell line, consisting of only a single HIV integration site. Thus, the activities of the validated LPAs and/or LRA synergizers may depend on the HIV proviral integration site specific to this cell line.

To further scrutinize the discovered LPAs and LRA synergizers for their dependency on HIV integration sites, the compounds were subject to additional latency reversal assays on two distinct Jurkat latency model cell lines, JLat 8.4 and JLat 6.3^35^. Due to compound availability, 2 out of the 18 hit compounds were not included in the experiment (B02 and E02, Table 1). The results indicated that all 14 tested LRA synergizers continued to synergize with TNF in JLat 8.4 and 6.3 cells (orange bars, Fig. 4A and 4B). Due to cell viability concerns, the LPA B15 remains inconclusive on its integration site dependency (grey bars, Fig. 4A and 4B). Overall, the results confirms that the compounds from the initial validation experiment reliably exert their effects on HIV latency reactivation across all three HIV proviral integration sites tested.

**Figure 4.**
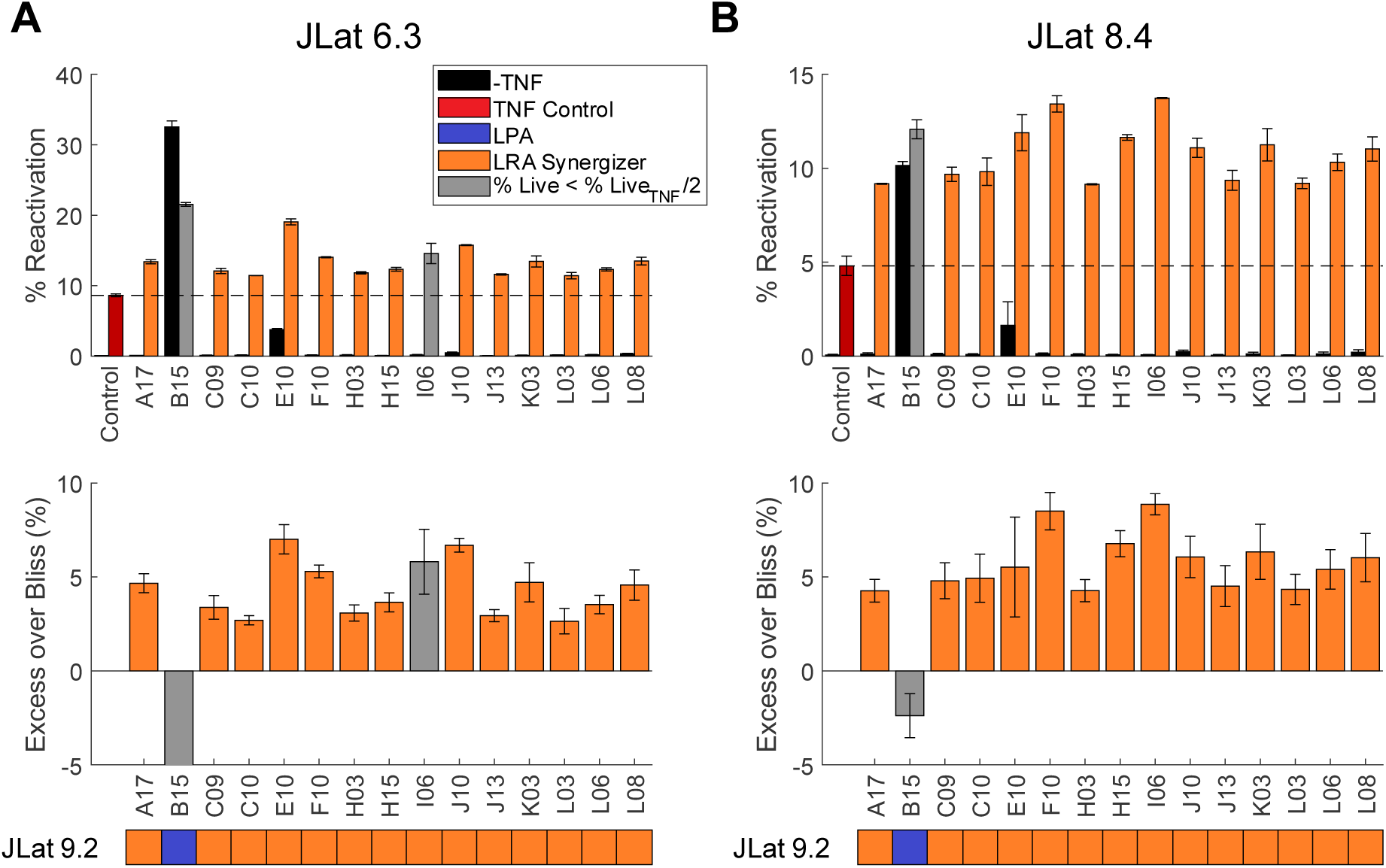
The effects of the experimentally validated latency modulating compounds are observed in two additional HIV integration sites. Additional latency reversal assays were conducted on JLat 6.3 and JLat 8.4 to investigate whether the effects of the latency modulating compounds depend on the integration site specific to JLat 9.2. **(A)** The results from JLat 6.3 agreed with those from JLat 9.2. Compounds B15 and I06 resulted in significant cytotoxicity and were inconclusive in this experiment. **(B)** The results from JLat 6.3 also agreed with those from JLat 9.2. Compound B15 again displayed significant cytotoxicity and remained inconclusive, but compound I06 did not show significant cytotoxicity and agreed with its performance on JLat 9.2. Horizontal dashed lines indicate the reactivation percentages of the TNF-only controls. See Fig. S6 for full results of the assays.

## Discussion

HIV is characterized by noisy and excitable cell-fate decisions, where gene expression fluctuations play a critical role in deciding whether an infected cell initiates viral production immediately upon viral infection, or develops into latency where the integrated provirus remains silent for months to years^15,43^. Single-cell time-lapse microscopy data of HIV gene expression dynamics can provide valuable insights into the effects of drug perturbations on cells, effectively capturing drug-target interactions. However, such large-scale experiments are time-and resource-intensive, and consequently, are not favorable in the pursuit of accelerated drug screening and discovery. In this study, seeking to incorporate HIV gene expression dynamics while circumventing these drawbacks, we employed a VAE model to learn latent representations of 1,806 unique compounds from an experimental dataset^17^ that captures unique features of both drug-induced HIV gene expression perturbations and the chemical structures of the drugs. These latent features enable us to infer novel HIV latency modulators by identifying drugs that share similar latent space representations with known latency modulators. However, our VAE model requires the input of time-dependent fluorescence noise trajectories, which are infeasible to obtain for the 175,000 compounds in our *in silico* screening. We circumvented this problem by concatenating time-series fluorescence noise trajectories of confirmed HIV latency modulators from the training set with the chemical structures of the compounds during the inference stage. This allowed us to investigate novel molecules under the premise that they elicit gene expression perturbations known to modulate latency in HIV and scored the likelihood of each compound to modulate HIV latency by comparing the similarity of their latent representations to those of known latency modulators. We further experimentally verified the top 113 predicted compound using a pre-established HIV latency cell model JLat 9.2, which resulted in the confirmation of 2 LPAs and 16 LRA synergizers^35^. However, by combining the chemical structures of screening compounds with the noise trajectories of confirmed HIV latency modulators, our computational approach may introduce some bias, as it assumes similar fluorescence noise trajectories across all latency modulators, potentially limiting the accuracy of predictions for novel candidates. Despite the potential bias in our model, and the sparsity of positive training data points (∼0.6% LPAs, ∼1.9% LRA synergizers), our ML-guided screening method was able to yield experimental hit rates (∼1.8% for LPAs, ∼13.3% for LRA synergizers) much higher than those reported by traditional drug screening modalities that rely on experimental techniques alone (<2%, commonly 0.01% - 0.14%)^44^, providing an efficient alternative to labor-and cost-intensive large-scale high-throughput screenings. Due to the varying nature of HIV integration sites in the host cell genome^37,38^, we examined whether the observed effects of the hit compounds is dependent on the integration site specific to JLat 9.2. We performed additional assays on JLat 6.3 and 8.4 with the hit compounds and confirmed that their effects are present for all three integration sites tested.

For the experimental validation of the ML predictions, we used a previously reported HIV latency model cell line JLat^35^ created using the leukemia cell line Jurkat. Although the JLat cell lines are well established and have been widely used for HIV research, they do not accurately recapitulate latent HIV infection *in vivo*. Further studies on the hit compounds must be conducted using primary HIV latency cell models and/or patient samples to examine their safety and effectiveness. On the other hand, HIV proviral integration into host genome is known to be semi-random^37^, and we examined three different integration sites using different JLat cell lines. Our results indicate that the effects of our hit compounds are consistent on all three integration sites tested. However, this does not rule out potential latent HIV integration sites that may be resilient to the hit compounds. Further validations of the compounds with many more integration sites, potentially utilizing polyclonal cell lines, are needed to provide a more comprehensive picture.

In summary, our study utilized advanced machine learning techniques by combining a variational autoencoder and a chemical language model to extract meaningful low-dimensional representations of the noise in HIV gene expression when exposed to a library of 1,806 compounds from a previous study^17^. These representations were then combined with the chemical fingerprint of a library of ∼175,000 compounds in an *in silico* screen for additional HIV latency modulators. The machine learning model utilized a previously discovered correlation between HIV noise modulators and HIV latency modulators, and accelerated the screening process by circumventing the need for experimental determination of noise modulating compounds through time-lapse fluorescence microscopy experiments. Despite the lack of positive training data, our model yielded a final experimental hit rate of 15.9%, where 18 out of the 113 predicted compounds were confirmed to be HIV latency modulators. The continued use of advanced machine learning methods similar to our method may provide additional leverage for accelerating drug discovery, especially in fields of study rich with available large-scale datasets.

## Materials and Methods

### Machine Learning Model Training and Inference

#### Training Stage

A total of 1,838 independent time-lapse fluorescence microscopy measurements were used in this study. It includes 1,806 compounds from a diverse library of chemicals, 72 TNF-treated positive controls, 72 alsterpaullone (Alp)-treated negative controls, and 144 untreated neutral controls, generated in Lu et al^17^, adding up to 2,094 independent measurements. A quality control preprocessing, identical to that reported in Lu et al^17^, was applied to this dataset prior to training, which excluded 268 measurements, bringing the total number down to 1,826. An additional 12 independent time-lapse fluorescence microscopy measurements on four LPAs (PX12, 60 μM; tiopronin, 4 mM; NSC400938, 10 μM; NSC 155706, 10 μM) reported in Lu et al, as well as TNF (10 ng/ml) and untreated controls^17^ were conducted in duplicate (n = 2) and combined with the aforementioned dataset, totaling 1,838 independent measurements. For each measurement, the noise in HIV gene expression is extracted from raw fluorescence microscopy data collected over an 18-hour period and was used as part of the training data. Molecular structures were encoded using CDDD^22^, a sequence-to-sequence autoencoder trained on chemically diverse molecules from the PubChem and ZINC datasets^32,33^ to translate between different SMILES representations of the same molecules. It mapped the SMILES strings for each of the 1,806 compounds to a continuous 512-dimensional latent vector which captured chemical semantics and structure. These embeddings, combined with the time series fluorescence data, were input into a Variational Autoencoder (VAE) with two hidden layers of 256 and 32 neurons in both the encoder and decoder. The VAE, trained for 200 epochs with a learning rate of 0.001 and batch size of 128, generated a 4-dimensional latent space to represent each compound. The VAE loss function was calculated as follows:

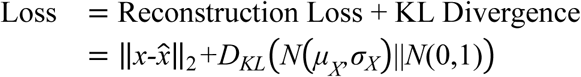

#### Inference Stage

The trained VAE model was then employed to screen potential HIV latency modulators within a collection of large compound libraries at the University of Illinois Urbana-Champaign (UIUC) High Throughput Screening Facility (HTSF)^23^, totaling to about 175,000 unique compounds. For each compound in the collection, CDDD embeddings were generated and concatenated with fluorescence noise data from confirmed LPA/LRA synergizer compounds in the training set. This combined input was passed through the VAE to project each compound into the latent space. Pairwise distances between the latent representations of known LPA/LRA synergizers and the library compounds were calculated, enabling the ranking of candidates. The top 100 compounds for both LPA and LRA synergizer categories were selected for experimental validation. After accounting for overlaps between the two categories, a total of 113 unique compounds were experimentally verified. This was expected since the model focusses on identifying HIV latency modulators, instead of trying to separate latency promoters and reversal synergizers in the latent space.

### Experimental Methods

#### Cell Culture

Jurkat cell lines (JLat 9.2, 8.4 and 6.3) were cultured in Corning RPMI 1640 with L-glutamine and phenol red. 10% of fetal bovine serum (FBS) and 1% of penicillin-streptomycin were supplemented to the culture media (final concentration: 100 U/ml penicillin, 100 ng/ml streptomycin). Cell cultures were started from-80°C backup vials at 1 ml total volume in T25 tissue culture treated flasks. The total volume was doubled when reaching high confluency (indicated by media color change from red/pink to yellow) up to 8 ml, and was then transferred and expanded to 24 ml in T75 tissue culture treated flasks. Cell cultures were passaged twice a week and maintained at 24 ml, with a dilution ratio of 1:3 (cell culture: fresh media) for each passage.

#### Latency Reversal Assays

Latency reversal assays (Fig. 3 and 4) were performed using flow cytometry. One day before drug treatments, the concentrations of the cell cultures of interest were measured using hemocytometers. The cultures were subsequently diluted to about 800,000 cells/mL with fresh media, while maintaining constant volume. On the day of experiment, cells were deposited into 96-well V-bottom plates, with 200uL of cell culture per well. Wells were then treated with the compounds of interest, with and without the latency reversing agent (LRA) tumor necrosis factor (TNF). All treatment conditions were performed in duplicate (n = 2). After treatment, samples were incubated at 37°C and 5% CO_2_ for 24 hours. The JLat 9.2 samples were then stained with propidium iodide (PI, 1 mg/ml, 1uL per well) before flow cytometry measurements, while the JLat 8.3 and 6.4 were measured directly. A BD LSR Fortessa Flow Cytometry Analyzer was used for the JLat 9.2 latency reversal assays, while a BD Symphony A1 Flow Cytometry Analyzer was used for the JLat 8.3 and 6.4 assays. For all latency reversal assays, a BD High Throughput Sampler was used for sample intake from the 96-well V-bottom plates. Forward scatter, side scatter, GFP fluorescence (488 nm excitation, 530/30 emission bandpass filter) and PI fluorescence (561 nm excitation, 610/20 emission bandpass filter, JLat 9.2 only) were measured. Cell viability (% live) is quantified using a combination of forward scatter, side scatter and PI fluorescence (JLat 9.2 only). PI staining was omitted for JLat 8.3 and 6.4 because it was sufficient to quantify JLat viability through forward scatter and side scatter alone (Fig. S5). Percentage of reactivated cells (% reactivated) is quantified using GFP fluorescence. For each sample, at least 15,000 live cells were collected.

#### Excess over Bliss (EoB) Score Calculation

To quantify the interaction between the compounds of interest and LRAs, we utilized a model reported in previous literature, called the Bliss independence model, to calculate the Excess over Bliss (EoB) score^36^. Let *f*_LRA_ and *f*_D_ denote the reactivation percentage of the LRA-only sample and the compound of interest-only sample, and *f*_LRA+D_ denote the reactivation percentage of the sample treated with both LRA and compound of interest. Then the EoB is defined as the difference between the measured combined effect of the two compounds and the expected combined effect. The expected combined effect is calculated assuming independence between the LRA and the compound of interest. The exact formula is as follows:

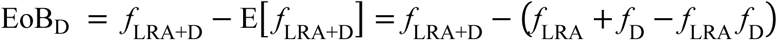

For compounds with EoB > 0, they are synergistic with the LRA; for those with EoB < 0, they are antagonistic with the LRA. The magnitudes (absolute values) of the EoB scores indicate the strength of synergism or antagonism. To exclude potential false positives arising from the high number of compounds tested, we did not use EoB = 0 as a threshold. Instead, a threshold on the magnitude of EoB was implemented for each day of experiment. The threshold is set as the median of the magnitude of all EoB scores obtained on the same day.

Since all samples were measured in duplicate, an error bar is calculated for each EoB score. To further reduce false positives, a compound is only considered as an LRA synergizer if its error bar is fully above the threshold, i.e. its minimum possible EoB score is above the threshold. A compound is only considered as an LPA if its error bar is fully below the threshold, i.e. its maximum possible EoB score is below the threshold.

## Data Availability

The image and signal datasets generated and analyzed for the 1,806 compounds and control treatments from the original 2021 screen are available at this University of Illinois at Urbana-Champaign Illinois Data Bank deposit: https://doi.org/10.13012/B2IDB-8103861_V1 (65).

The code for the feature engineering, model training, inference, and data visualization in this study can be found at this Shukla Group GitHub repository: https://github.com/ShuklaGroup/HIV-Latency-VAE.

## Acknowledgements

This work was supported by National Institutes of Health grants R35GM142745 and R21AI167693 awarded to D.S, and the Chemistry-Biology Interface Research Training Program (T32-GM070421) and Samuel W. Parr Fellowship to J.H. We thank the Roy J. Carver Biotechnology center, Cytometry and Microscopy to Omics Facility (RRID:SCR_025272 https://coremarketplace.org/?FacilityID=2727) for providing expertise in flow cytometry for our project.

## References

1. Douek, D. C., Roederer, M. & Koup, R. A. Emerging Concepts in the Immunopathogenesis of AIDS. Annu. Rev. Med. 60, 471–484 (2009).

2. Powell, M. K. et al. Opportunistic Infections in HIV-Infected Patients Differ Strongly in Frequencies and Spectra between Patients with Low CD4+ Cell Counts Examined Postmortem and Compensated Patients Examined Antemortem Irrespective of the HAART Era. PLoS ONE 11, e0162704 (2016).

3. Barré-Sinoussi, F. et al. Isolation of a T-Lymphotropic Retrovirus from a Patient at Risk for Acquired Immune Deficiency Syndrome (AIDS). Science 220, 868–871 (1983).

4. Wagner, T. A. et al. Proliferation of cells with HIV integrated into cancer genes contributes to persistent infection. Science 345, 570–573 (2014).

5. Maldarelli, F. et al. Specific HIV integration sites are linked to clonal expansion and persistence of infected cells. Science 345, 179–183 (2014).

6. Siliciano, J. D. et al. Long-term follow-up studies confirm the stability of the latent reservoir for HIV-1 in resting CD4+ T cells. Nat Med 9, 727–728 (2003).

7. Yeh, Y.-H. J. & Ho, Y.-C. Shock-and-kill versus block-and-lock: Targeting the fluctuating and heterogeneous HIV-1 gene expression. Proc. Natl. Acad. Sci. U.S.A. 118, e2103692118 (2021).

8. Janssens, J., et al. Mechanisms and efficacy of small molecule latency-promoting agents to inhibit HIV reactivation ex vivo. JCI Insight 9, e183084 (2024).

9. Debrabander, Q. et al. The efficacy and tolerability of latency-reversing agents in reactivating the HIV-1 reservoir in clinical studies: a systematic review. Journal of Virus Eradication 9, 100342 (2023).

10. Yates, K. B. et al. Epigenetic scars of CD8+ T cell exhaustion persist after cure of chronic infection in humans. Nat Immunol 22, 1020–1029 (2021).

11. Rutishauser, R. L., et al. TCF-1 regulates HIV-specific CD8+ T cell expansion capacity. JCI Insight 6, e136648 (2021).

12. Kessing, C. F. et al. In Vivo Suppression of HIV Rebound by Didehydro-Cortistatin A, a “Block-and-Lock” Strategy for HIV-1 Treatment. Cell Reports 21, 600–611 (2017).

13. Singh, A. & Weinberger, L. S. Stochastic gene expression as a molecular switch for viral latency. Current Opinion in Microbiology 12, 460–466 (2009).

14. Singh, A., Razooky, B., Cox, C. D., Simpson, M. L. & Weinberger, L. S. Transcriptional Bursting from the HIV-1 Promoter Is a Significant Source of Stochastic Noise in HIV-1 Gene Expression. Biophysical Journal 98, L32–L34 (2010).

15. Weinberger, L. S., Burnett, J. C., Toettcher, J. E., Arkin, A. P. & Schaffer, D. V. Stochastic Gene Expression in a Lentiviral Positive-Feedback Loop: HIV-1 Tat Fluctuations Drive Phenotypic Diversity. Cell 122, 169–182 (2005).

16. Dar, R. D., Hosmane, N. N., Arkin, M. R., Siliciano, R. F. & Weinberger, L. S. Screening for noise in gene expression identifies drug synergies. Science 344, 1392–1396 (2014).

17. Lu, Y. et al. Screening for gene expression fluctuations reveals latency-promoting agents of HIV. Proc. Natl. Acad. Sci. U.S.A. 118, e2012191118 (2021).

18. Diederik, P. K. & Max, W. An Introduction to Variational Autoencoders. Foundations and Trends® in Machine Learning 12, 307–392 (2019).

19. Goodfellow, I. et al. Generative adversarial networks. Commun. ACM 63, 139–144 (2020).

20. Grønbech, C. H. et al. scVAE: variational auto-encoders for single-cell gene expression data. Bioinformatics 36, 4415–4422 (2020).

21. Yoon, J., Jarrett, D. & van der Schaar, M. Time-series Generative Adversarial Networks. in Advances in Neural Information Processing Systems (eds Wallach, H. et al.) vol. 32 (Curran Associates, Inc., 2019).

22. Winter, R., Montanari, F., Noé, F. & Clevert, D.-A. Learning continuous and data-driven molecular descriptors by translating equivalent chemical representations. Chem. Sci. 10, 1692–1701 (2019).

23. Krishnamurthy, V. Compound Collections. Compound Collections | School of Chemical Science https://scs.illinois.edu/resources/cores-scs-research-and-service-facilities/high-throughput-screening-facility/compound.

24. Sukuru, S. C. K. et al. Plate-Based Diversity Selection Based on Empirical HTS Data to Enhance the Number of Hits and Their Chemical Diversity. SLAS Discovery 14, 690–699 (2009).

25. Zhu, W. et al. Identification of SARS-CoV-2 3CL Protease Inhibitors by a Quantitative High-Throughput Screening. ACS Pharmacol. Transl. Sci. 3, 1008–1016 (2020).

26. Dreiman, G. H. S., Bictash, M., Fish, P. V., Griffin, L. & Svensson, F. Changing the HTS Paradigm: AI-Driven Iterative Screening for Hit Finding. SLAS Discovery 26, 257–262 (2021).

27. Hansen, M. M. K. et al. A Post-Transcriptional Feedback Mechanism for Noise Suppression and Fate Stabilization. Cell 173, 1609–1621.e15 (2018).

28. Bohn-Wippert, K. et al. Cell Size-Based Decision-Making of a Viral Gene Circuit. Cell Reports 25, 3844–3857.e5 (2018).

29. Ross, J. et al. Large-scale chemical language representations capture molecular structure and properties. Nat Mach Intell 4, 1256–1264 (2022).

30. Jaeger, S., Fulle, S. & Turk, S. Mol2vec: Unsupervised Machine Learning Approach with Chemical Intuition. J. Chem. Inf. Model. 58, 27–35 (2018).

31. Weininger, D. SMILES, a chemical language and information system. 1. Introduction to methodology and encoding rules. Journal of chemical information and computer sciences 28, 31–36 (1988).

32. Irwin, J. J. et al. ZINC20—A Free Ultralarge-Scale Chemical Database for Ligand Discovery. J. Chem. Inf. Model. 60, 6065–6073 (2020).

33. Kim, S. et al. PubChem 2025 update. Nucleic Acids Research 53, D1516–D1525 (2025).

34. Simidjievski, N. et al. Variational Autoencoders for Cancer Data Integration: Design Principles and Computational Practice. Front. Genet. 10, 1205 (2019).

35. Jordan, A. HIV reproducibly establishes a latent infection after acute infection of T cells in vitro. The EMBO Journal 22, 1868–1877 (2003).

36. Bliss, C. I. THE TOXICITY OF POISONS APPLIED JOINTLY1. Annals of Applied Biology 26, 585–615 (1939).

37. Wang, G. P., Ciuffi, A., Leipzig, J., Berry, C. C. & Bushman, F. D. HIV integration site selection: Analysis by massively parallel pyrosequencing reveals association with epigenetic modifications. Genome Res. 17, 1186–1194 (2007).

38. Craigie, R. & Bushman, F. D. HIV DNA Integration. Cold Spring Harbor Perspectives in Medicine 2, a006890–a006890 (2012).

39. Jordan, A. The site of HIV-1 integration in the human genome determines basal transcriptional activity and response to Tat transactivation. The EMBO Journal 20, 1726–1738 (2001).

40. Rezaei, S. D. & Cameron, P. U. Human Immunodeficiency Virus (HIV)-1 Integration Sites in Viral Latency. Curr HIV/AIDS Rep 12, 88–96 (2015).

41. Einkauf, K. B. et al. Intact HIV-1 proviruses accumulate at distinct chromosomal positions during prolonged antiretroviral therapy. Journal of Clinical Investigation 129, 988–998 (2019).

42. Jiang, C. et al. Distinct viral reservoirs in individuals with spontaneous control of HIV-1. Nature 585, 261–267 (2020).

43. Pal, S. & Dhar, R. Living in a noisy world—origins of gene expression noise and its impact on cellular decision-making. FEBS Letters 598, 1673–1691 (2024).

44. Zhu, T. et al. Hit Identification and Optimization in Virtual Screening: Practical Recommendations Based on a Critical Literature Analysis: Miniperspective. J. Med. Chem. 56, 6560–6572 (2013).

