## Supplementary Information for "Variational Autoencoder-enabled High-throughput Drug Screening for HIV Latency Modulators predicted through Noise in Gene Expression"

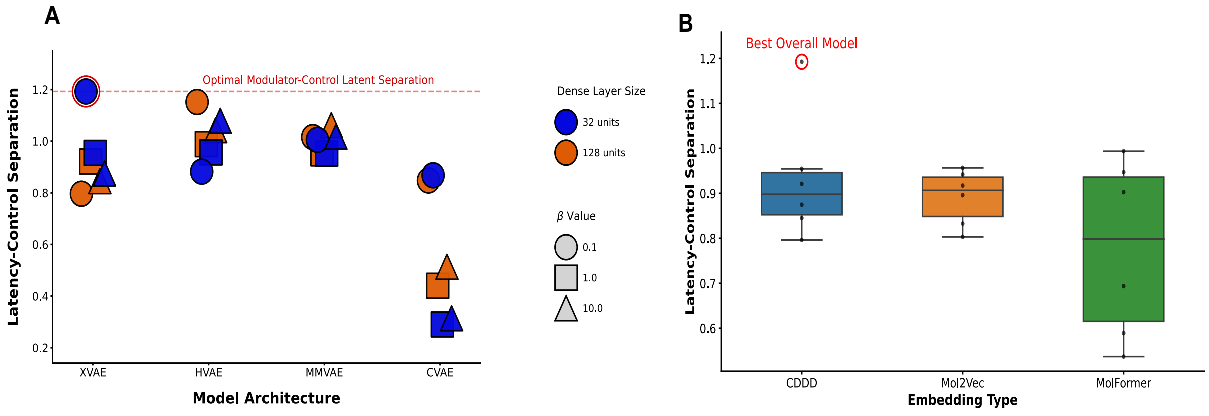


**Figure S1. VAE architecture and embedding selection based on latent separation of validated modulators from controls. (A)** We compared the mean pairwise distances between validated HIV latency modulators and non-modulating control compounds within the latent space of each model. Greater separation indicates improved discrimination and learned correlation between proximity and modulatory activity. The XVAE model with a latent layer size of 4, regularization strength (β) of 0.1 and dense hidden layer size of 32 was selected through this ablation. **(B)** After selecting this XVAE model, we selected CDDD as the best input embedding since it gave the largest latency-control separation by far, despite Mol2vec having a slightly higher median separation distance.

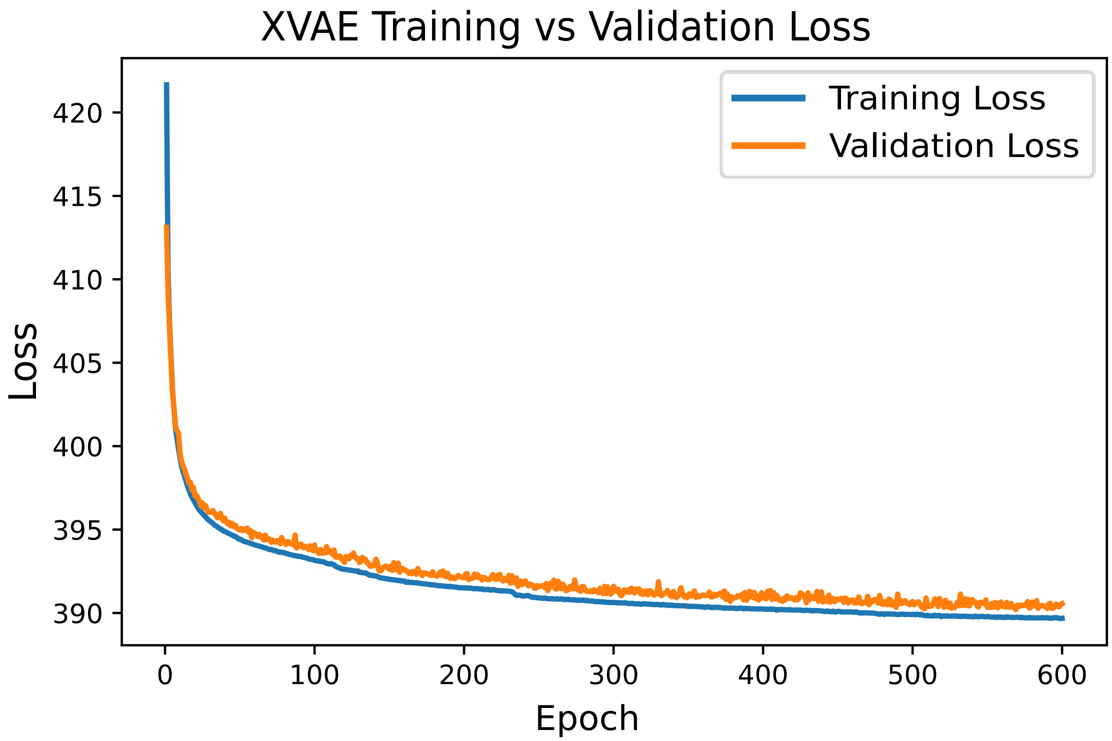


**Figure S2. Training and validation loss vs training epochs.** The VAE model learns important patterns from training data without overfitting, as shown by both training loss on seen data and validation loss on unseen data steadily decreasing and plateauing after 370 epochs of training.

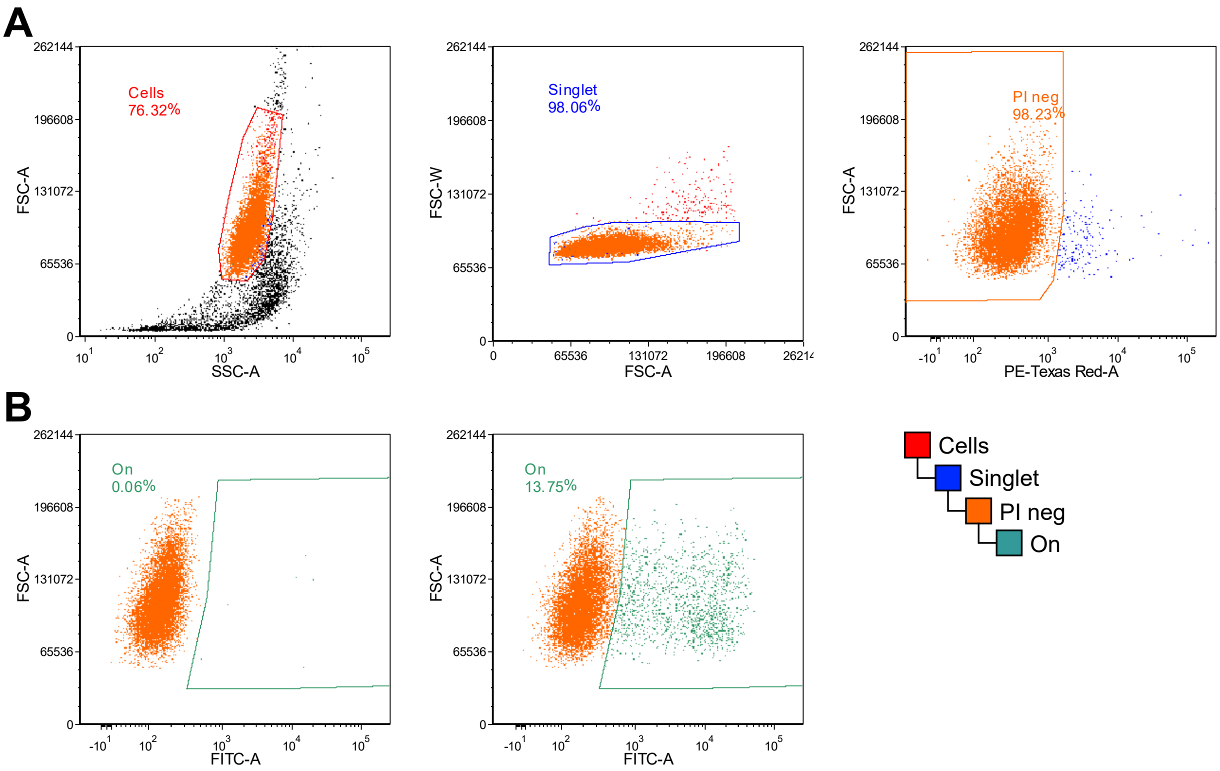


**Figure S3. Gating strategy for JLat latency reversal assays. (A)** The gates used to determine valid events collected from the flow cytometers. The "Cells" gate (left) contains the highest density of events plotted on forward scatter area (FSC-A) vs. side scatter area (SSC-A). The "Singlet" gate (middle) is a subgate to the "Cells" gate that includes events within a small range of forward scatter width (FSC-W) to remove cell clumps. The "PI neg" gate (right) is a subgate to the "Singlet" gate where cells low on propidium iodide (PI) fluorescence (PE-Texas Red-A) are included. PI is a cell stain that selectively stains dead cells. **(B)** Gating strategy used to determine HIV reactivation. The "On" gate is a subgate of the "PI neg" gate and includes cells that are high in GFP fluorescence (FITC-A). Left panel shows an example result obtained from an untreated sample, while the right panel shows the result from a TNF-treated sample.

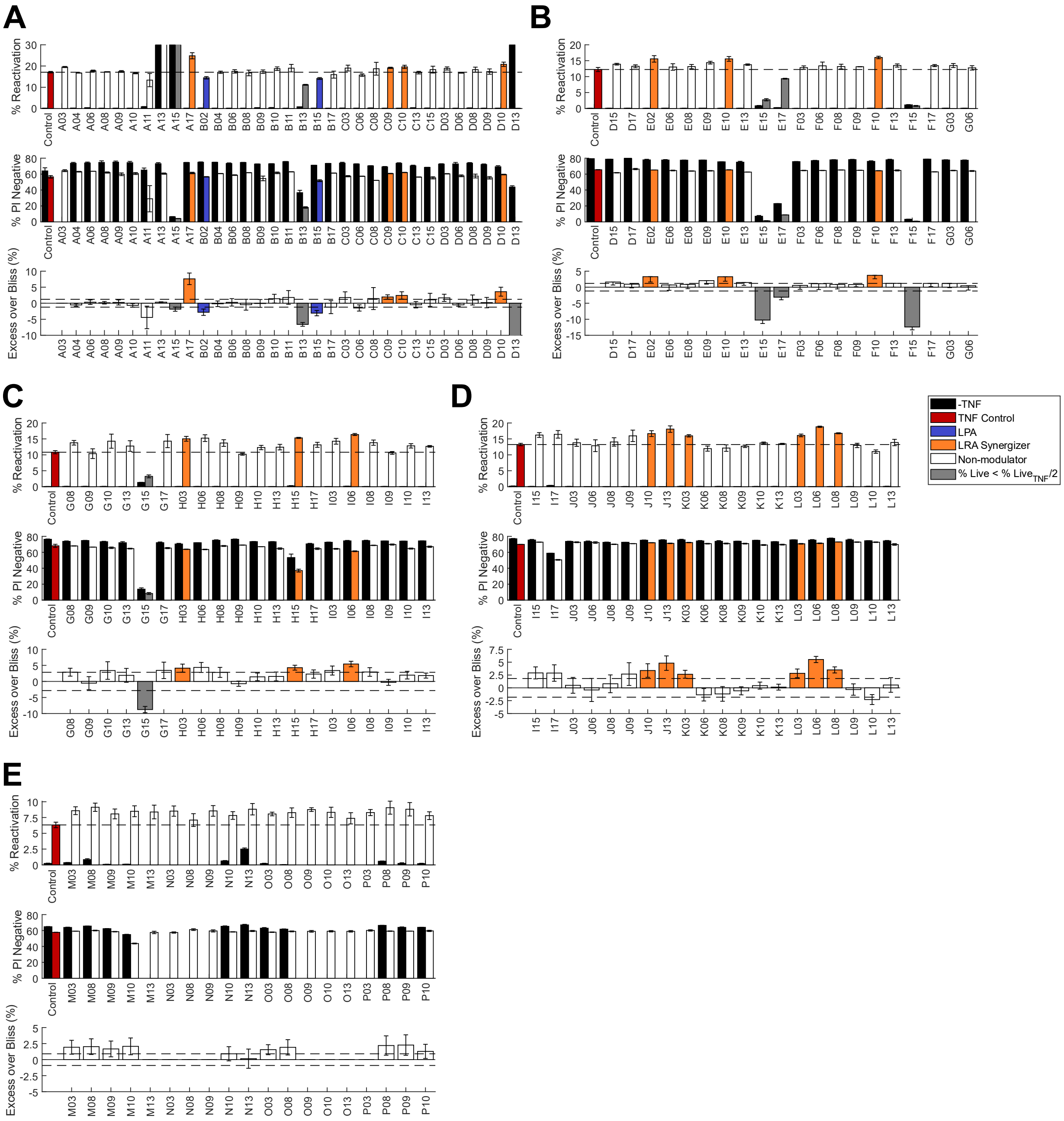


**Figure S4. Full result of the JLat 9.2 latency reversal assay on machine learning predicted compounds.** Panels **(A) – (E)** corresponds to the results from experiment batches 1 – 5, each carried out on a separate day due to throughput constraints. "% Reactivation" refers to the percentage ratio of events in the "On" gate over the events in the "PI neg" gate (Fig. S3). "% PI Negative" refers to the percentage ratio of events in the "PI neg" gate over all events collected in the sample (Fig. S3). See methods section for Excess over Bliss (EoB) score calculation. Due to limited supply, the EoB score of a few compounds were not obtained due to missing –TNF samples. Notable examples include A03 in panel (A), and M13, N03, N08, N09, O09, O10, O13, P03 in panel (E). Horizontal dashed lines indicate the reactivation percentage of the TNF only controls.

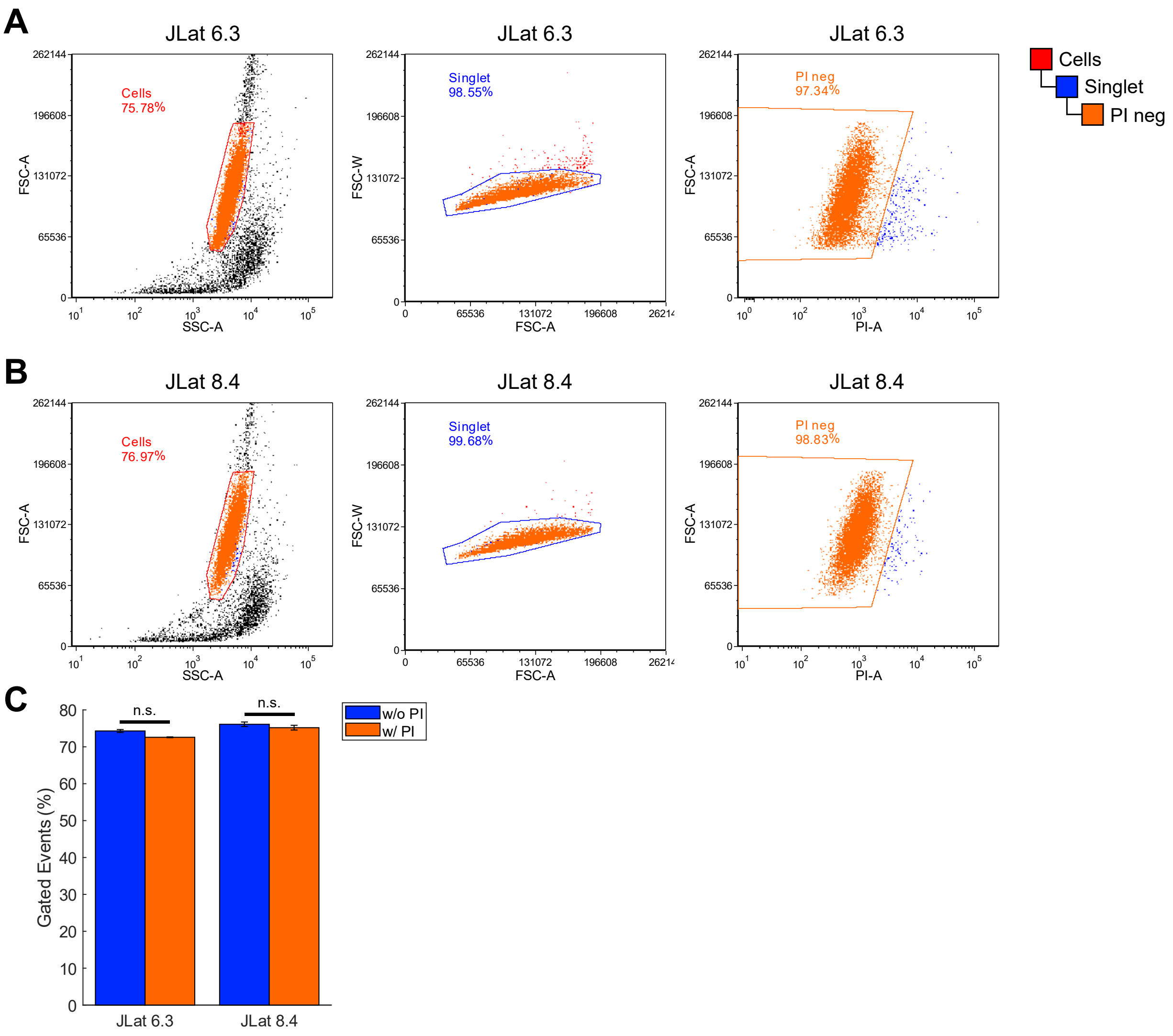


**Figure S5. Comparison of cell viability quantification with and without PI staining for JLat 6.3 and 8.4.** Additional control experiments were carried out with JLat 6.3 and 8.4 cells stained with PI to examine whether the use of cell viability staining result in statistically significant shift in the gating results. **(A)** Results from JLat 6.3. **(B)** Results from JLat 8.4. Gate hierarchy is shown to the right side of the figure. **(C)** Comparison between gating results with and without PI gating. Blue bars represent results obtained from only the "Cells" gate and the "Singlet" gate. Orange bars represent results obtained from all 3 gates. The effect sizes of the two set of samples are relatively small. In addition, two-sample *t*-tests were carried out and both JLat 6.3 and JLat 8.4 showed no statistical significance ($p > 0.05$) between the gating results. For JLat 6.3, $p = 0.0518$. For JLat 8.4, $p = 0.4098$.

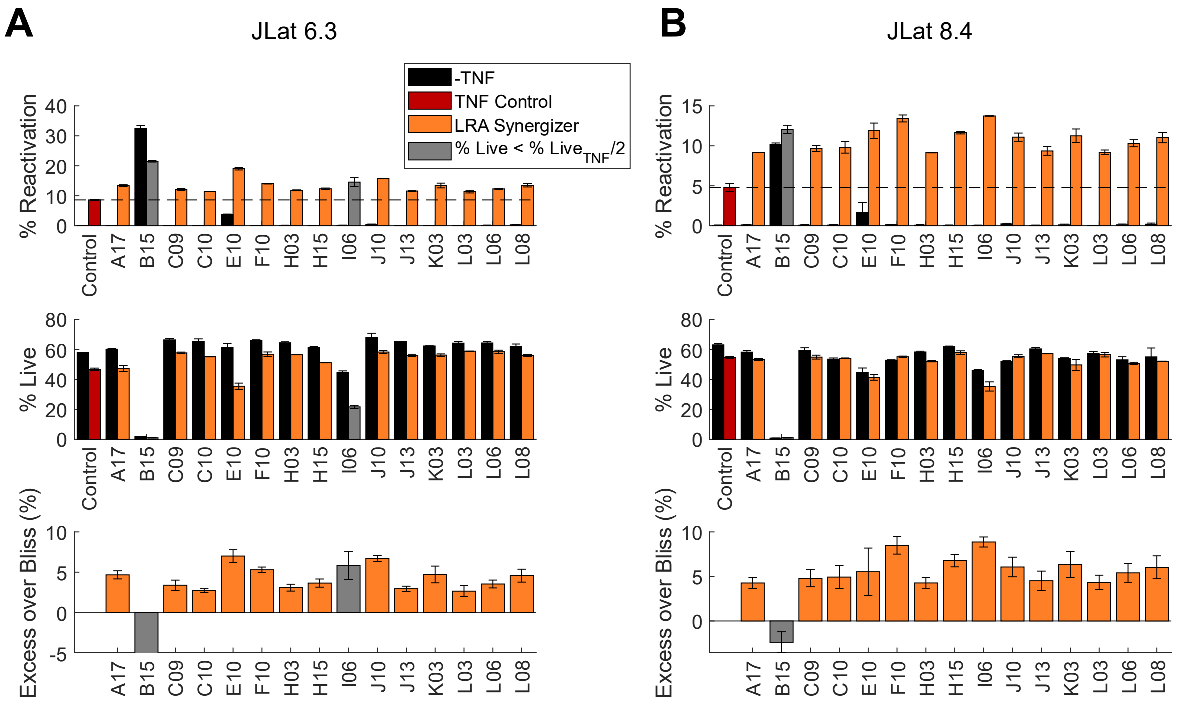


**Figure S6. Full result of the JLat 6.3 and 8.4 latency reversal assay on latency modulator hits from JLat 9.2 assay.** Panels **(A)** and **(B)** show the results of latency reversal assay, based on JLat 6.3 and 8.4 respectively, using compounds listed in Table 1. Compounds B02 and E02 were unavailable from suppliers and was not investigated. The first row shows the percentage of cells reactivating out of latency, quantified by GFP fluorescence. The second row shows the percentage of live cells, quantified by forward and side scatter. See Fig. S3 and S5 for gating details. The third row shows the excess over Bliss (EoB) score of each compound. Horizontal lines indicate the reactivation percentages of the TNF-only controls.
